# Do divergent mitochondrial backgrounds alter the fate of nuclear alleles during hybridization and admixture?

**DOI:** 10.64898/2026.11.04.764285

**Authors:** Leah J. Darwin, Rebecca Z. Bachtel, Faye A. Lemieux, Yevgeniy Raynes, David M. Rand

## Abstract

Hybridization brings together genetic variation from different populations or species into admixed genomes. As selection sorts this variation, the fate of each allele is determined by its fitness effect in the new hybrid genomic background. Because mitochondrial genes coevolve with the nuclear genome, they may be especially sensitive to foreign nuclear backgrounds, making them potential “hot-spots” for hybrid incompatibilities. Here, we test whether selection against mitonuclear incompatibilities drives repeatable hybridization outcomes. Specifically, we experimentally evolved three sets of five populations of hybrid *Drosophila melanogaster*, each carrying the same nuclear chromosomes but paired with one of three increasingly divergent mitochondrial DNA backgrounds. We hypothesized that accumulated incompatibilities would result in different patterns of nuclear admixture among mitochondrial backgrounds. To test this, we tracked nuclear allele frequencies in all 15 populations by whole-genome sequencing at intermediate time points. After 25 generations, we observed that rapid genomic evolution produced a strikingly parallel mosaic of both nuclear ancestries, independent of mitochondrial background. Contrary to our hypothesis, we found no evidence that mitochondrial background shaped nuclear admixture, even for interspecific combinations. These results suggest that in species with large effective population sizes, efficient purifying selection limits the accumulation of mitonuclear incompatibilities, allowing selection on nuclear alleles to promote repeatable ancestry sorting immediately following hybridization.

## Introduction

Hybridization, the interbreeding of genetically distinct populations or species, is increasingly recognized as an important source of genetic variation, producing admixed individuals whose genomes are mosaics of both ancestries [Taylor and Larson, 2019, Moran et al., 2021]. Natural selection subsequently filters this variation, retaining advantageous alleles and purging deleterious or incompatible alleles from the genome [Juric et al., 2016, Matute et al., 2020, Langdon et al., 2022]. Notably, genomic outcomes across independent hybridization events may be repeatable as selection consistently acts on the same general pool of genetic variation, including those alleles intrinsically incompatible with each other [Schumer et al., 2018, Chaturvedi et al., 2020, Farleigh et al., 2026]. Here, we ask whether selection on incompatibilities between mitochondrial and nuclear genomes shapes repeatable hybridization outcomes.

Mitochondrial and nuclear genes involved in the oxidative phosphorylation (OXPHOS) system may be “hotspots” for intrinsic incompatibilities because they encode protein subunits that must function together for essential cellular function [Rand et al., 2004, Moran et al., 2021]. Given the central role of the OXPHOS system in producing cellular energy in eukaryotes and its broad effects on fitness [Gershoni et al., 2009], selection may be expected to strongly favor coordinated function between the two genomes, promoting mitonuclear coevolution. Additionally, the mitochondrial genome’s high mutation rate and reduced effective population size [Lynch et al., 2006, Lynch and Blanchard, 1998] may further drive coevolution through nuclear compensation, where the nuclear genome evolves to compensate for deleterious mutations in the mitochondrial genome [Rand et al., 2004, Osada and Akashi, 2012]. Mitonuclear incompatibilities are part of a broader class of barriers to hybridization, Bateson-Dobzhansky-Muller incompatibilities (BDMIs), in which alleles viable within each parental population become deleterious when expressed together in hybrids [Orr and Turelli, 2001]. BDMIs have been identified in a variety of systems [Zuellig and Sweigart, 2018, White et al., 2012, Schumer et al., 2014, Bikard et al., 2009] and are particularly abundant in OXPHOS genes [Meiklejohn et al., 2013,Lopez et al., 2021,Pereira et al., 2021, Moran et al., 2024, Robles et al., 2025]. In fact, mitonuclear incompatibilities are often suggested to play a disproportionately large role in the process of speciation [Hill, 2017, Sloan et al., 2017]. The mitonu-clear system thus presents a powerful framework for understanding how limits to hybridization emerge from disrupting coevolved molecular systems.

Here, we developed an experimental system to test the repeatability of admixture patterns following intraspecific hybridization. Three sets of five populations were initiated from a hybridization between two North American inbred laboratory-adapted strains of *Drosophila melanogaster*, where each set carried one of three increasingly divergent mitochondrial DNA (mtDNA) backgrounds: two from diverged *D. melanogaster* populations and one from a different species. We hypothesized that accumulated BDMIs in increasingly divergent mtDNAs would create differences in nuclear admixture patterns. To test this prediction, we measured the genomic changes of the 15 independently evolving populations using whole-genome sequencing at intermediate time points. After over 40 generations of experimental evolution under laboratory conditions, we instead found that rapid genomic evolution in the hybrids resulted in a strikingly parallel mosaic of both North American ancestries independent of the mtDNA background. Our findings suggest that in species with large effective population sizes like *Drosophila*, nuclear fitness differences may outweigh the fitness effects of mitonuclear incompatibilities due to efficient purifying selection acting on mtDNA haplotypes.

## Results

To test whether mitonuclear incompatibilities affect the repeatability of genome evolution following hybridization, we established replicate admixed populations that differed in their mitochondrial DNA (mtDNA) background. For each mtDNA background, we initiated 5 replicate laboratory populations by reciprocal F1 and F2 crosses between two common North American lines of *D. melanogaster* : OreR (Ore) collected from Oregon and Ral375 (Ral) collected from Raleigh, North Carolina (Figure 1). Before the establishment of hybrid populations, each mtDNA was introgressed onto the Ore and Ral nuclear backgrounds and homogenized across groups through round-robin crosses (see Methods). The three mitochondrial haplotypes were selected to represent increasing levels of divergence: a *D. melanogaster* mtDNA from a Beijing population (diverged ∼3k years ago from North American populations [Arguello et al., 2019]); a *D. melanogaster* mtDNA from a Zimbabwe population (diverged ∼12k years ago from North American populations [Arguello et al., 2019]); and an interspecific mtDNA from *D. yakuba* (diverged ∼8–10M years ago from *D. melanogaster* [Tamura, 2003]). The populations, consisting of ∼4300 individuals (see Figure S1 for all population size estimates), were maintained under standard laboratory conditions for over 40 generations with periodic whole-genome sequencing to quantify changes in nuclear allele frequencies over time.

**Figure 1:**
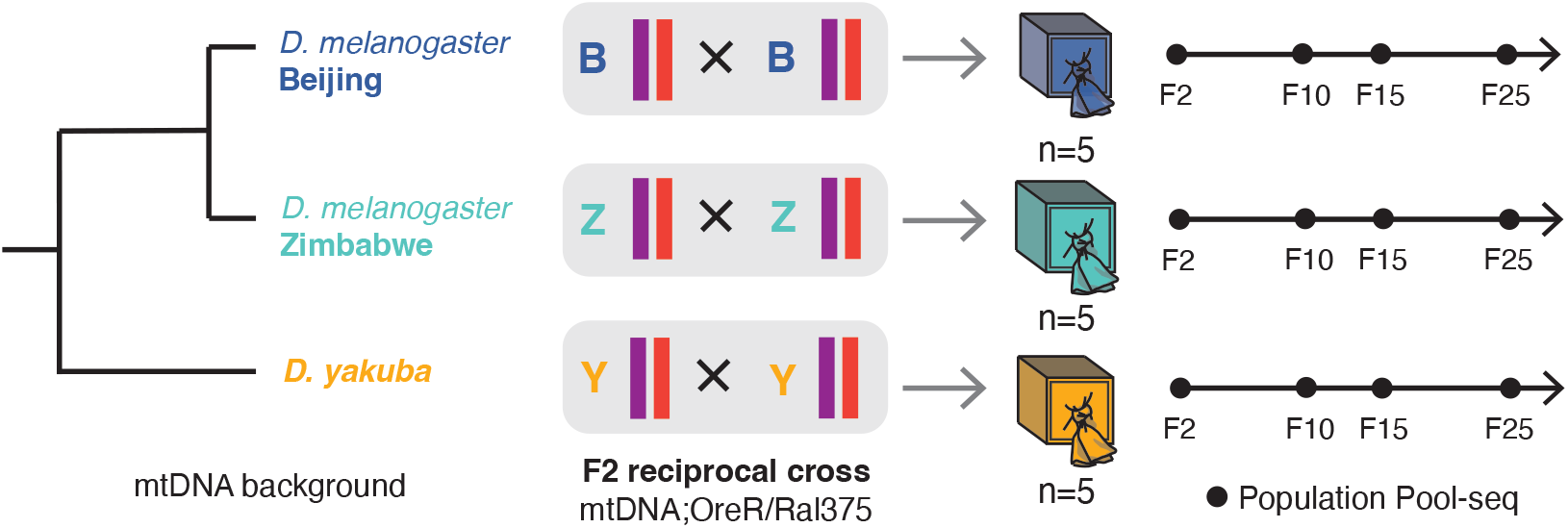
Experimental evolution of hybrid nuclear genomes across divergent mtDNA backgrounds. Fifteen experimental hybrid populations of *D. melanogaster* were experimentally evolved and differed only in their mitochondrial background (mtDNA; shown in phylogenetic tree). Each population was initiated at approximately 50/50 nuclear ancestry from an F2 reciprocal cross between F1 hybrids carrying the parental nuclear chromosomes OreR and Ral375. For each mtDNA background, *n* =5 populations were initiated and evolved under laboratory conditions. Pooled-sequencing (pool-seq) was performed at intermediate generations (shown with solid points).

### Laboratory evolution produces consistent phenotypic outcomes

We first tested whether divergent mitochondrial backgrounds produced differences in two fitness-related phenotypes in the evolved populations: female fecundity and development time. We measured female fecundity at generation F42 by counting the total offspring for each of 45 mated females per population. We found no significant effect of the population’s mtDNA background on female fecundity (*p* = 0.759, Tables S1–S2, Figure 2a). At generation F25, we measured the egg-to-pupa development time. We found no significant effect of the mtDNA background on development time (*p* = 0.233, Tables S1–Table S2, Figure 2b). After over 25 generations of experimental evolution, we observed no phenotypic fitness differences attributable to divergent mtDNA effects. This indicates that divergent mtDNA backgrounds imposed no measurable fitness cost in these hybrids, providing little phenotypic evidence for the mitonuclear incompatibilities we expected to shape admixture.

**Figure 2:**
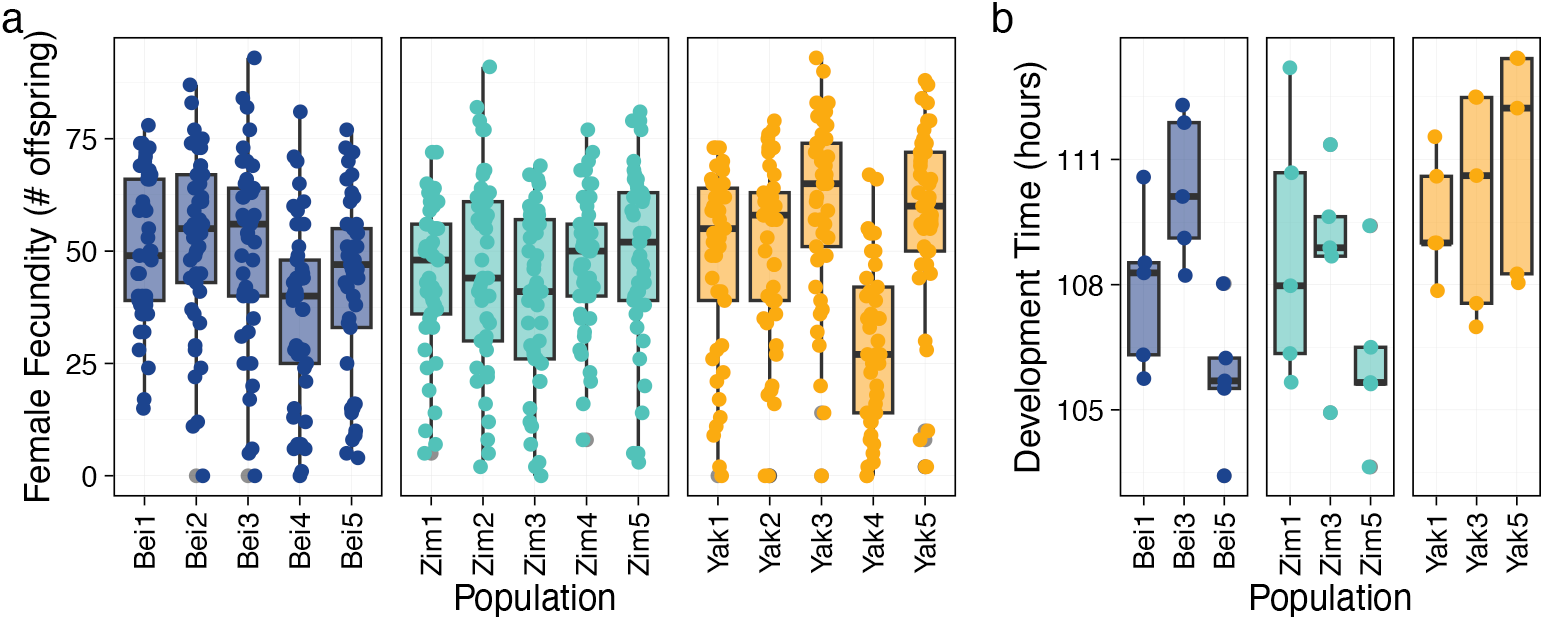
Phenotypic comparisons between evolved populations. **(a)** Boxplots show female fecundity at generation F42 for each replicate population. Each overlaid point represents the total offspring for a single mated female. **(b)** Boxplots show the measures of development time at F25 for samples of individuals from a subset of populations. Each overlaid point represents the average development time for a single density-controlled vial of 40 individuals.

### Rapid and highly parallel hybrid genome evolution

To test if admixture patterns differed between populations harboring different mtDNA genotypes, we performed pooled sequencing of all populations at intermediate time points (Figure 1b). After 25 generations of evolution, we identified 388,595 SNPs that changed significantly through time using a generalized linear model test (Equations 4–5, Figure 3a). This indicates that there was a rapid response to hybridization that spanned a substantial proportion of the genome. Across all 15 populations, the largest axis of variation in genome-wide allele frequencies reflected shared change over time rather than differences between populations, indicating parallel evolution (Figure 3b, Figure S2). To resolve whether this shared response reflected repeatable changes in nuclear ancestry, we quantified how the frequency of each nuclear ancestry (Ore/Ral) changed during the experiment using a previously described method [Linder et al., 2020]. At generation F2, Ore and Ral were present at approximately equal frequencies across the majority of chromosome 3 (Figure S3a). Chromosome 2 and the X chromosome appeared to be subject to segregation bias at F2, as the Ore genotype frequency was closer to 40% than the expected 50% (Figure S3a). However, there was no effect of mtDNA background on chromosomal segregation at generation F2 (genome-wide Kruskal-Wallis test, Figure S4). At generation F25, there were large deviations in the frequencies of nuclear ancestries from the initial starting frequencies on chromosomes 2, 3, and X (Figure 3c). These ancestry deviations were associated with the significant changes in SNP frequencies (Figure 3a,c). Most strikingly, the Ore genotype on chromosome arm 3L increased in parallel across all populations, rising from 50% to nearly 100% frequency (Figure 3c). We also observed very parallel evolution across the other autosomes but with less dramatic deviations from the starting frequencies (Figure 3c). The X chromosome showed a parallel decrease in Ore frequency spanning much of its length (Figure 3c). There was no statistically detectable effect of mtDNA background on the predicted Ore genotype frequencies at F25 (genome-wide Kruskal-Wallis test, Figure S4). This high degree of parallelism likely reflects selection on the shared variation between populations, with coordinated frequency shifts at physically adjacent sites indicating linked selection. In general, increases in both nuclear ancestries largely balanced each other across the genome, leaving the genome-wide ratio of ancestries close to 50:50 (e.g., 46% Ore, 54% Ral at generation F25). Together, the genetic similarity among mitochondrial backgrounds and comparable phenotypic fitness suggest that there was no detectable effect of mitonuclear incompatibilities accumulated with mtDNA divergence on the genomic patterns of admixture in these populations.

**Figure 3:**
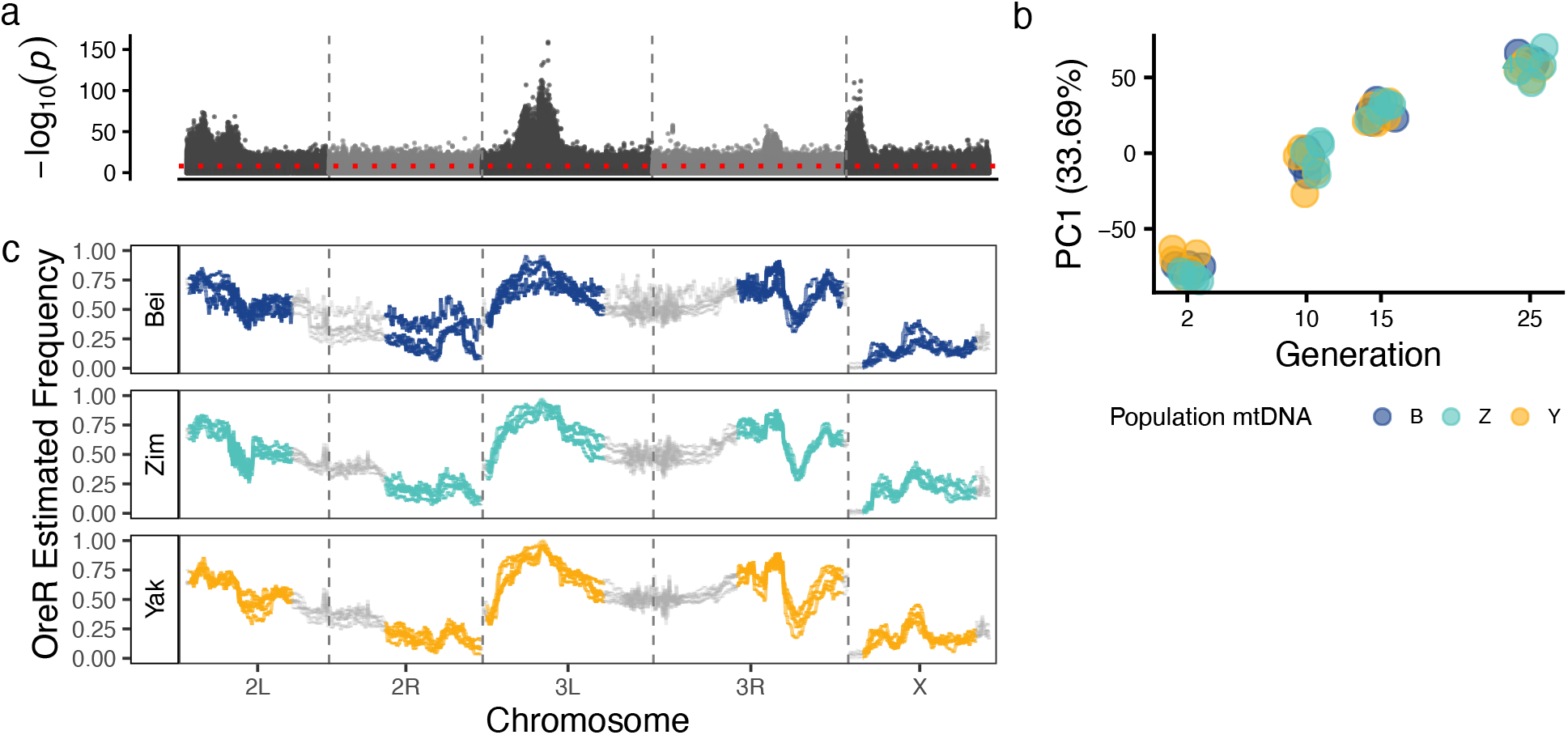
Parallel genomic hybrid evolution and ancestry sorting. **(a)** For each SNP, the − *log*_10_(*p*) value for a significance test of time is plotted for the autosomes and X chromosome using data from all populations. A quasibinomial generalized linear model was fit to each variant and the significance of generation time was tested using a likelihood ratio test. The dotted red line gives the Bonferroni-adjusted significance threshold. Significant SNPs indicate consistent changes in allele frequencies through time. **(b)** PCA analysis of genome-wide allele frequencies over time. The greatest axis of variation (PC1, 33.69%) is explained by shared changes in allele frequencies in all populations (replicate B3 excluded from analysis due to contamination). Each point is a replicate population colored by the population mtDNA (Beijing (B), Zimbabwe (Z), and *D. yakuba* (Y)). **(c)** For each mtDNA population type, the estimated frequency of the genotype OreR (Ore) over chromosomes 2, 3, and X are plotted. Populations are grouped into panels by mtDNA (Beijing (Bei), Zimbabwe (Zim), and *D. yakuba* (Yak)) and each line represents one of 5 population replicates per group. The alternative nuclear genotype, Ral375, is not shown as it is at frequency 1 − OreR. Centromeric and telomeric regions are masked out in grey as they do not resolve easily to a specific ancestry resulting in noise around 50% and are excluded from analysis (based on coordinates given in Table 3 of [Comeron et al., 2012] lifted over to the dm6 reference genome).

Because recombination has been suggested to shape the evolution of hybrid genomes [Schumer et al., 2018], we tested whether recombination rate was correlated with ancestry frequency in our populations. Since each chromosome arm had distinct patterns of the selected nuclear ancestry, we tested for correlations with recombination rate independently on each chromosome arm. Chromosome arms 3L and 3R exhibited elevated Ore ancestry (71% and 62%, respectively, Figure 3c) and a significant positive correlation between Ore frequency and recombination rate (3L: *ρ* = 0.333, *p <* 0.001; 3R: *ρ* = 0.209, *p* = 0.013, Figure S5, Table S3). Conversely, chromosome arm 2R was enriched for Ral ancestry (80%) and exhibited a negative correlation between Ore frequency and recombination rate (*ρ* = − 0.272, *p* = 0.003, Figure S5, Table S3), corresponding to a positive correlation between recombination rate and Ral frequency. Chromosome arm 2L did not show a significant correlation (Figure S5, Table S3). Overall, when a given ancestry had a greater proportion on a given chromosome arm, the relationship between its local frequency and recombination was positive, consistent with selection being more effective in regions of high recombination. The X chromosome, which is hemizygous, retained the lowest proportion of OreR ancestry of any chromosome (19%; Figure 3c) and showed no relationship between OreR frequency and recombination rate (Figure S5, Table S3), suggesting strong and uniform intolerance for admixture across this chromosome.

### Statistical mtDNA effects are coincidental, not causal

In addition to analyzing differences in ancestry across mtDNA backgrounds, we performed SNP-based tests for selection using the time series data (Figure 1b). Following [Wiberg et al., 2017], we fit a quasibinomial generalized linear model for each SNP and tested for time-dependent effects of mtDNA background using a likelihood ratio test of the time ×mtDNA interaction effect (Equations 4, 6). We identified a group of physically adjacent SNPs on chromosome arm 2L with significant time × mtDNA interaction effects, indicating allele frequency changes that differed with mtDNA background (Figure 4a). We found that populations with a Zim mtDNA background had consistently elevated allele frequency changes from F2 to F25 (| F2 − F25 |) for significant SNPs in this region (Figure 4b). Zim mtDNA population-specific effects were also found to explain 4.48% of the total genetic variation along principal component 3 (Figure S2).

**Figure 4:**
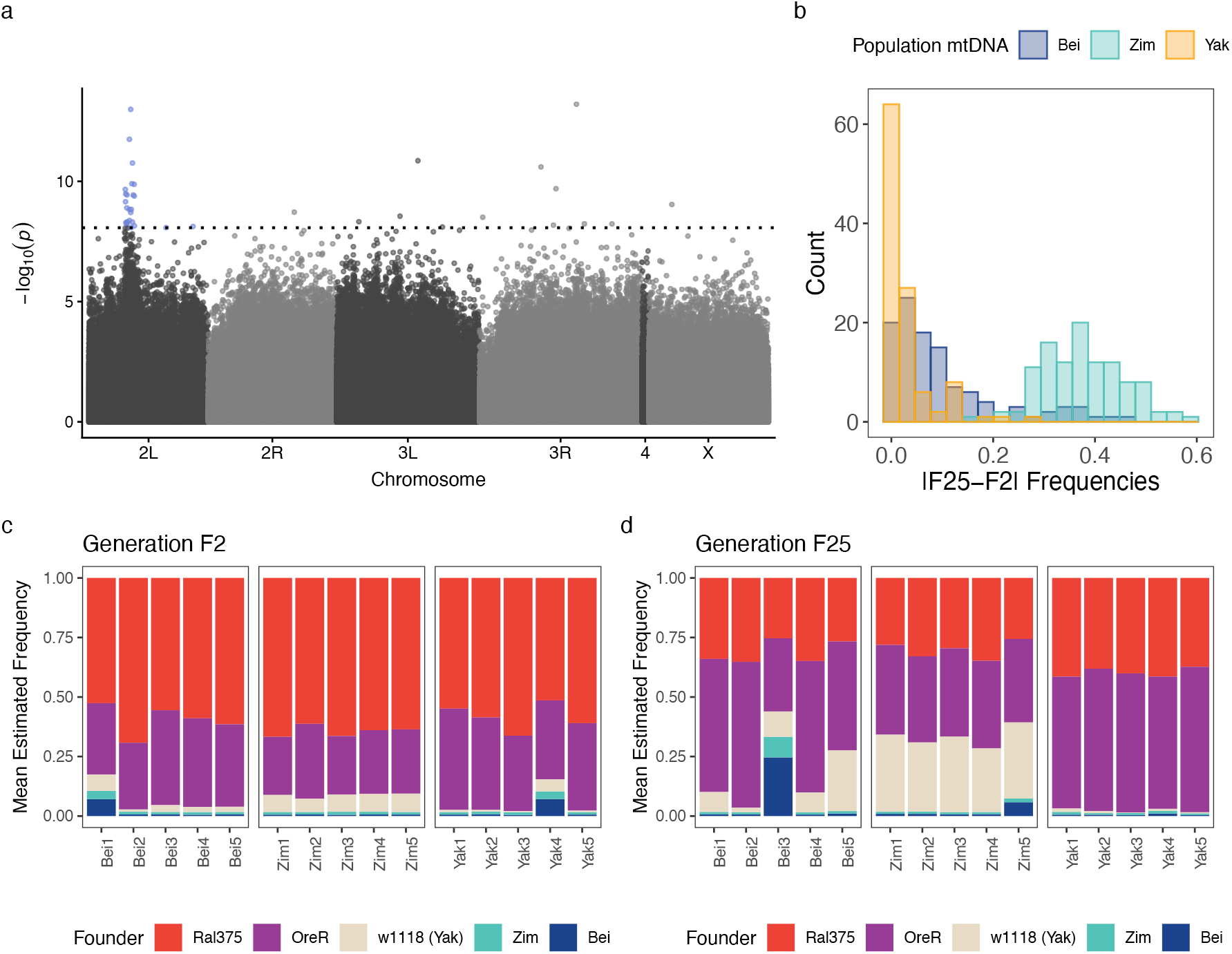
SNP-based tests of mtDNA effects. **(a)** Manhattan plot of *p*-values for likelihood ratio test on time×mtDNA interaction effect. The dotted line shows the Bonferroni-adjusted significance threshold. On chromosome arm 2L, a region consistent with a signature of linked selection is highlighted. **(b)** Histograms of allele frequency changes between generation F2 and F25 (|F2 − F25|) for significant SNPs (highlighted, panel a). All populations for each mtDNA type: Beijing (Bei), Zimbabwe (Zim), and *D. yakuba* (Yak) are combined into a single distribution. For the region covered by the significant SNPs (highlighted in panel (a)), average estimates of all nuclear DNA backgrounds used in the backcrossing scheme and the round-robin crosses are shown for each population (mtDNA1–5) and generations **(c)** F2 and **(d)** F25.

Because Ore and Ral ancestry did not explain the Zim mtDNA-specific allele frequency changes, we tested whether contamination from additional backgrounds contributed to genetic differentiation. Each mtDNA originally resided on a different nuclear background (*mtDNA;nucDNA*; *Bei;Bei, Zim;Zim, Yak;w*^1118^) before being introgressed onto the shared Ore and Ral nuclear backgrounds through 10 generations of repeated backcrossing, followed by round-robin randomization. Despite this level of nuclear homogenization, it remains possible that the initial nuclear chromosome type contributed a small amount of genetic variation to our founder lines by chance. Consistent with this hypothesis, all five Zim mtDNA populations showed a consistent increase in *w*^1118^ ancestry in the region with Zim mtDNA dependent SNP frequency changes (Figure 4d), reflecting transfer of the *w*^1118^ background between mtDNA lines during round-robin crossing (see Methods).

Several lines of evidence suggest that the increase in *w*^1118^ frequency among Zim mtDNA populations does not reflect fitness effects of mitonuclear incompatibilities. First, one population with a Beijing mtDNA background (Bei5) showed a similar increase in *w*^1118^ frequency in this region by F25 (Figure 4d). Second, all five populations with the Zim mtDNA had a significantly higher estimated starting F2 frequency of the *w*^1118^ background within this region (*p* = 0.0023, via Wilcoxon rank-sum test; Figure 4c). This is further supported by the observation that Zim populations were already differentiated on PC3 at F2 (Figure S2). Increases in the *w*^1118^ background were not specific to the Zim mtDNA populations, and *w*^1118^ likely had higher starting frequencies in the Zim mtDNA populations by chance. Other SNPs with significant time×mtDNA interaction effects not in the highlighted region of chromosome 2L (Figure 4a) were associated with very subtle differences in allele frequency changes among mtDNA backgrounds (Figure S6). Because mitonuclear incompatibilities would likely cause substantial, linked changes in allele frequencies, it is more likely that these additional significant SNPs are false positives for selection on mitonuclear incompatibilities. Based on these observations, the most parsimonious interpretation is that the observed statistical associations with mtDNA background do not reflect the influence of mitonuclear incompatibilities on evolution following hybridization.

Over the course of the experiment, we identified additional instances of contamination from other backgrounds as well as other technical issues associated with the long-term maintenance of populations. Populations that stood out as outliers from the other replicates were identified with PCA analysis of genome-wide allele frequencies. On PC1 and PC2, the population replicate Bei3 was an outlier only at generation F25 (Figure S2), due to a severe population bottleneck introduced by a technical issue (e.g., Figure 4d). Additionally, on PC2, four additional populations were outliers at F2 or F10 (Bei1, Bei2, Yak4, Yak5; Figure S2). Again, we suspect that this is due to rare haplotypes from other backgrounds (e.g., Figure 4c). These populations were not outliers at F15 or F25, indicating that the contaminating background was either eventually lost or maintained at undetectable frequencies (*<* 1*/*2*n* = 0.25% given our sample size of 200 diploids). When these time points were excluded, reanalysis of the data did not change our qualitative conclusions (Figure S7), implying that any possible contamination did not meaningfully affect our results. More broadly, these observations highlight the difficulty of achieving complete homogenization of nuclear backgrounds through standard laboratory methods.

## Discussion

In this study we directly tested the repeatability of intraspecific hybrid genome evolution across three increasingly divergent mtDNA backgrounds. To do so, we experimentally evolved 15 outbred populations of *D. melanogaster* for over 40 generations under laboratory conditions and quantified allele frequency changes through time. Our major finding was that nuclear genome evolution in hybrid populations remained highly repeatable across mitochondrial backgrounds, even for mtDNAs derived from a species that diverged ∼8–10M years ago.

While we did not find evidence that BDMIs arising from interactions with the mitochondrial genome shaped hybrid evolution, the unique distribution of allele frequency changes on chromosome arm 2L in the Zimbabwe mtDNA populations suggests a possible association. However, the elevated frequency of *w*^1118^ at generation F2 in the Zimbabwe populations was also present in the Bei3 and Bei5 populations, which stood out in the PCA analyses. For this pattern to reflect an effect of Zim mtDNA, the *w*^1118^ alleles would have needed a fitness advantage in Zim mtDNA background and a separate, unique fitness advantage in the Bei3 and Bei5 populations. A more parsimonious explanation is that the *w*^1118^ background had a stochastically elevated frequency in the founder Zim mtDNA line after round-robin randomization, which was then propagated into all five replicate populations and subsequently realized its *w*^1118^-specific fitness effects in those particular contexts. For these reasons, the simplest interpretation of our results is that different mtDNA backgrounds have limited, if any, impact on the fate of competing nuclear genomic regions in this experiment.

More broadly, this result does not support the hypothesis that mitonuclear interactions disproportionately affect hybridization [Gershoni et al., 2009, Burton and Barreto, 2012, Hill, 2017]. We posit that observations of influential mitonuclear BDMIs in other systems such as copepods [Pereira et al., 2021] and swordtail fishes [Moran et al., 2024,Robles et al., 2025] are not an inevitable consequence of mtDNA divergence. The formation of mitonuclear BDMIs may be promoted by certain species- or population-specific factors, such as high mtDNA mutation rates, small effective population sizes, population subdivision, or intense functional constraints [Sloan et al., 2017]. The relatively large effective population sizes of *D. melanogaster* (∼ 1.5M, [Duchen et al., 2013]) and *D. yakuba* (∼ 6.5M, [Sawyer and Hartl, 1992]) may have facilitated efficient purifying selection against deleterious mtDNA variants, leaving little opportunity for mitonuclear compensatory coevolution. Consistent with this, dN/dS ratios for nuclear genes that encode proteins that contact mtDNA-encoded proteins do not exceed 1 in bivalves [Maeda et al., 2021] and mammals [Weaver et al., 2022], whereas values greater than 1 would be expected if positive selection were acting on compensatory nuclear loci. The lack of a relationship between mtDNA divergence and its impacts on hybridization observed here highlights the conditions under which mitonuclear BDMIs are expected to arise.

Several features specific to our experimental system may have limited our power to detect mitonuclear interaction effects that are well documented elsewhere. Mitochondrial interactions are known to affect a wide range of traits [Immonen et al., 2016, Camus and Inwongwan, 2023, Nguyen et al., 2023, Nguyen et al., 2025], though these effects are often small relative to nuclear and environmental effects [Mossman et al., 2016, Darwin et al., 2025]. The latter study confirms that there are highly significant differences between the OreR and Ral375 nuclear genomes for important performance traits [Darwin et al., 2025]. The relatively small populations (∼ 4300 individuals) used in this experiment may therefore have been insufficient for selection on the fitness effects of mitonuclear interactions to overcome stochastic allele frequency changes due to drift alone. A complementary study to the one presented here tracked competing mtDNA haplotypes on different constant nuclear backgrounds in outcrossing *Drosophila* populations [Mossman et al., 2019]. Mitonuclear effects on mtDNA frequency changes were generally inconsistent, providing little evidence that nuclear background predictably influenced mtDNA dynamics, consistent with what we observed here. In contrast, a study that quantified changes in nuclear allele frequencies on constant mtDNA backgrounds in asexually evolving yeast found significant effects of mtDNA background on nuclear allele frequency trajectories [Biot-Pelletier et al., 2023]. The differences in our findings may be explained by larger population sizes or more generations of evolution, both of which increase the opportunity for weak selection to leave a detectable signal. Notably, neither of these studies initiated populations with hybrid nuclear backgrounds, a novel contribution that distinguishes the present work. The results presented here suggest that the strong selective pressure on nuclear alleles immediately following hybridization drives rapid ancestry sorting, rendering any potential, comparatively weak selective effects of mitonuclear interactions undetectable in admixture outcomes.

In the absence of strong mitonuclear BDMIs, there are other mechanisms that may explain the observed levels of parallelism. First, selection was likely acting primarily on shared nuclear variation across the three sets of five replicate populations. We used a round-robin crossing scheme to distribute any line-specific variation across the founding lines before starting the experiment. Consequently, the standing variation among the populations at the beginning of the experiment was very similar, although it is virtually impossible to ensure that all populations have identical initial allele frequencies at all loci. Populations that share a significant portion of standing genetic variation and are exposed to the same environments are more likely to evolve in parallel [Schlö tterer, 2023]. Second, the nuclear backgrounds used in this study were highly inbred and harbored more deleterious alleles than lines sampled from natural populations. As a result, parallel changes may be attributable to the purging of universally deleterious alleles. Other studies of parallel evolution in hybrids show that deleterious “hybrid” load from smaller populations is repeatably purged [Juric et al., 2016, Matute et al., 2020]. Our findings suggest a similar mechanism, but without a fitness asymmetry between the hybridizing populations, we did not observe a regression towards a particular parental population (e.g., the OreR alleles were advantageous on chromosome 2L but deleterious on the X). We also cannot rule out incompatibilities or other epistatic effects between the two nuclear genomes as contributors to this parallelism. Pooled-sequencing data lack the resolution to test for such effects directly, which is a limitation of this study. Despite this limitation, the observed parallelism suggests that the genomic effects of hybridization between closely related populations may be highly repeatable even in the absence of a change in the environment.

In conclusion, our results could not explicitly assign causal effects of mitochondrial background on nuclear allele fate during hybridization and admixture in this system. Moreover, the idea that mitonuclear BDMIs disproportionately affect hybridization broadly across species, or that mitonuclear coevolution is the functional basis of discordant patterns of mtDNA and nuclear loci in natural populations, should be met with considerable caution. Rather, mitochondrial DNA may simply be one genomic compartment that can accumulate incompatibilities when purifying selection is rendered inefficient by small effective population sizes, low recombination, high mutation rates, or strong functional constraint. Because these same conditions can affect other regions of the genome, future work on the predictability of admixture following hybridization may consider genetic regions that share these “mitochondrial-like” properties as a more general class of barriers to gene flow.

## Methods

### Population cage protocol

#### Genetic cross of founder lines

The founder lines were established in two phases: (1) genetically placing each mitochondrial DNA (mtDNA) haplotype onto two common nuclear backgrounds and (2) intermixing heterogeneity in the similar nuclear backgrounds across the mtDNA types with round-robin crosses (Figure S8). We selected three mtDNA haplotypes, two *D. melanogaster* mtDNAs from Beijing and Zimbabwe (Bei11, ZW142 [Grenier et al., 2015]), and one from

*D. yakuba* (Yak). The mitonuclear constructions were performed as described in [Darwin et al., 2025]. We will use the notation *mtDNA;nucDNA* to describe the mitonuclear constructions. Briefly, virgin females from each of the original lines carrying these mtDNAs: *Bei11;Bei11, ZW142;ZW142*, and *Yak;w* ^1118^ (interspecific construction originally described in [Ma et al., 2013, Spierer et al., 2021]) were crossed to males from each of two *D. melanogaster* lines: OreR (*Ore;Ore*) and Ral375 (*Ral;Ral*, also known as DGRP375), over 10 generations of recurrent backcrossing. While backcrossing eliminated the majority of the original background variation (theoretically 1/2^10^, or 0.0976% of the nuclear genome after 10 generations of backcrossing remains), we wanted any remaining heterogeneity among these lines to be evenly distributed across all lines so as not to confound the effect of the different mtDNA backgrounds. To accomplish this, we performed six generations of “round-robin” crosses within each nuclear background group. At each generation, virgin females from each backcross line were crossed to males from the other two lines that shared a nuclear genotype but differed by *mtDNA* genotype. Table S4 gives the initial three generations of these crosses for brevity.

#### Initiation of population cages

The 15 total population cages were initiated from crosses between the founder lines. For each mtDNA type, there were two “mixed” nuclear backgrounds each for OreR (Ore) and Ral375 (Ral) from the round-robin crossing process (four total nuclear backgrounds, Table S4). For each mtDNA type, we then performed a reciprocal F1 cross between the nuclear backgrounds (*mtDNA;Ore×mtDNA;Ral, mtDNA;Ral×mtDNA;Ore*). A total of 20 females and 20 males from each of the four nuclear backgrounds were used in five replicate crosses and the crosses were brooded to increase yield. The theoretical expectation is that the off-spring of these crosses will all share the same mtDNA background but will be heterozygous for Ral375 and OreR (*mtDNA*;*Ral/Ore, mtDNA*;*Ore/Ral* ). To ensure similar starting allele frequencies across mtDNA backgrounds, we additionally set up crosses between the F1 lines by performing an additional reciprocal cross (*mtDNA*;*Ral/Ore×mtDNA*;*Ore/Ral, mtDNA*;*Ore/Ral×mtDNA*;*Ral/Ore*). Both crosses were brooded, and each of the five population cages per mtDNA type was initiated with four standard fly rearing bottles, one from each direction of the cross and from each of the two broods. The choice of five replicate populations per mtDNA background was based on a balance of sequencing costs and the assumption that if a genomic region showed a shared pattern of allele frequency change in all five replicates of one mtDNA background relative to the others, it would provide some statistical support for a role of that mtDNA in driving nuclear allele evolution (e.g., 1*/*(0.5^5^)= 0.03, or unlikely by chance alone).

#### Population cage maintenance

Fly populations were maintained in clear acrylic cages (20.5 cm w x 14.5 cm d x 17.5 cm h) on a standard lab diet (10.4 g yellow cornmeal, 22 g sucrose, 1.8 g agar, 5 g SAF yeast, 0.9 g tegosept in 4.5 mL 95% ethanol, cooked in 200 mL of deionized water; individual batches were scaled up maintaining the same ratios). Six standard fly rearing bottles (30 mL of food per bottle) were in each population cage. Cages were maintained in a 25^*°*^ C incubator with a 12 hour light cycle. Every two weeks, six fresh bottles of food were placed inside the cages and flies were given 24 hours to lay eggs on the fresh food substrate. After 24 hrs, bottles were cleared of adult flies and maintained in an incubator at 25^*°*^C. Adults from the previous generation were collected by first placing the cages in a -20^*°*^C freezer and then moving the individuals to a -80^*°*^C freezer for long-term storage, which ensured non-overlapping generations. After one week, the bottles with offspring for the new generation were placed back in cages, beginning the cycle again.

#### Census population size estimates

To estimate the census population size, adult flies were frozen inside the cages and funneled into pre-weighed 25 ml centrifuge tubes following the 24 hour long egg collection at every generation. Flies were then weighed as a proxy for census population size, assuming one animal weighs 1 mg (from grand mean [Darwin et al., 2025]). Weight data were collected from F9 to F27. We observed that the population sizes oscillated, with gradual increases followed by rapid declines, which we attribute to an observed increase in larval density. Due to this observation, we modeled the estimated census population size (*N* ) as a function of generation number (Gen) using a linear harmonic regression of the form:

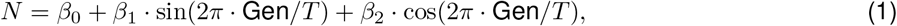

where *T* is the period of oscillation. The optimal period *T* was selected by searching a grid of candidate values (2 ≤*T ≤*18) and choosing the value that maximized the leave-one-out cross-validated *R*^2^. The mean population size was then estimated as the mean of the fitted curve over the observed generation range.

### Phenotypic assays and analysis

#### Development time assay

Three replicate cages from each mtDNA type (B1,B3,B5; Z1,Z3,Z5; Y1,Y3,Y5) were selected for the development time assay. After removing bottles of the outgoing F25 generation from the cages, grape juice agar plates smeared with yeast paste were placed inside the selected cages and flies were allowed to lay eggs for six hours. Plates were then removed from the cages, and eggs were picked from the plates and deposited in vials of food. Each vial received 40 eggs; there were 5 vials per cage. On the fourth day following the egg lay, vials were checked systematically every four hours between 8 AM and 8 PM. All pupae (pupal stage ≥P1 as defined by [Chyb and Gompel, 2013]) were counted, recorded, and marked on the vial to prevent double-counting. This continued for three days, stopping pupariation had largely ceased.

To test for an effect of the mitochondrial background common to a set of populations, we used the following mixed-effect linear model:

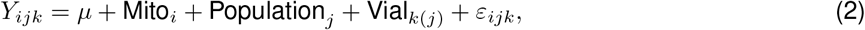

where *Y* is the development time of an individual, Mito is the fixed effect of the mtDNA background, Population is a random effect of the population from which individuals were sampled, and Vial is a random effect nested within Population that captures the shared environment of individuals that happened to be grown in the same vial. The significance of Mito was assessed using an F-test with Satterthwaite-approximated degrees of freedom. Models were implemented using the R (v4.5.2) packages lme4 (v2.0.1) and lmerTest (v3.2.1).

#### Fecundity assay

Fecundity was measured for a sample of 45 females from each population. During the population cycle of F42, when bottles were typically placed inside cages after a week of development, two bottles for each cage were left capped. Following seven days of further development and eclosion time, the capped bottles were removed from the cages and the adult flies were moved into a new bottle with fresh food. Separating the adults from other stages of development ensured that the flies used for the fecundity assay (1) were mature and (2) were mated. The flies were left in new bottles for three days to further ensure mating success. Individual females were then quickly distributed to 45 replicate vials per cage, limiting CO2 exposure to less than five mins. Flies were then placed into an incubator (25°C, 12 hour light cycle). Females were left to lay eggs for four days, after which the females were removed from vials. Any dead females were recorded. After ten days of development, the newly eclosed flies were counted.

To test for the effect of mtDNA background, we used the following mixed-effect linear model:

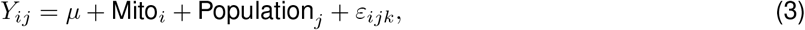

where *Y* is the number of offspring for each female, Mito is the fixed effect of the mtDNA background, and Population is the random effect of the population from which the females were sampled. The significance of Mito was assessed using an F-test with Satterthwaite-approximated degrees of freedom. Models were implemented using the R (v4.5.2) packages lme4 (v2.0.1) and lmerTest (v3.2.1).

### DNA sequencing and analysis

#### DNA extraction and sequencing

At generations F2 (starting generation), F10, F15, and F25, a random sample of 200 female individuals was selected from all 15 populations for pooled-sequencing. Animals were retrieved from storage in the -80^*°*^C freezer and sorted by sex on ice blocks. DNA was extracted from homogenized pools of whole animals using a chloroform/phenol-based protocol [Kapun et al., 2020]. The library preparation and sequencing were performed by Azenta/GENEWIZ and libraries were sequenced to ∼50x coverage using 150 bp paired-end reads.

#### Processing of DNA sequencing data

Raw sequencing data quality was assessed by Azenta/GENEWIZ, with additional manual verification using FastQC (v0.12.1) on both pre- and post-filtered reads. Reads failing to meet a minimum quality score of 20 or a minimum length of 50 bp were excluded from downstream analysis [Mallard et al., 2018]. The filtered reads were then aligned in paired-end mode to the D. melanogaster reference genome (FlyBase release 6.32) using BWA-MEM (v0.7.17) [Li and Durbin, 2009]. Following alignment, reads were converted to the pool-seq synchronized allele count (sync) format using PoPoolation2 [Kofler et al., 2011], applying a minimum mapping quality threshold of 20.

#### Ancestry Frequency Estimation

To estimate the frequencies of each parental genotype from the pooled-sequencing data, we first called variants from the previously sequenced inbred lines. Short reads from all of the lines were obtained from NCBI (PR-JNA1377152 [Darwin et al., 2025] for OreR and Ral375, *w*^1118^ data pending publication, and PRJNA268111 for Beijing and Zimbabwe). Reads were aligned and quality controlled using the same procedure as described in the section “Processing of DNA sequencing data.” We used a standard GATK (v4.5.0) pipeline to call variants from all of the inbred lines using base recalibration with a reference panel of known variants from the DGRP freeze 2 data [Huang et al., 2014]. Final variant filtering was applied according to the recommended GATK hard-filtering parameters [Van der Auwera et al., 2013]. The resulting VCF file was converted to tabular format using BCFtools (v1.21) query and a custom awk script, encoding the reference allele as 1, the alternative allele as 0, and heterozygous calls as missing, which were subsequently not used for predicting ancestry. We then used an adapted version of the fly XQTL script *haplotyper*.*limSolve* [Linder et al., 2020] to estimate the most likely proportion of each ancestry given the observed allele frequencies using a sliding window size of 100kb with a 10kb step size. We set the Gaussian weighting scheme such that the 500 SNPs closest to the window center accounted for 50% of the total weight. To identify windows that had uncertain ancestry predictions, we implemented a bootstrap procedure that resampled the SNPs in the window (with replacement) and predicted the ancestry proportions (*n* = 200 bootstraps, per window, per sample). Windows that had a standard deviation *>* 0.01 (*±* 1% frequency uncertainty) for either founder were subsequently excluded from analysis. These highly uncertain windows tended to localize to chromosome centromeres, consistent with the highly repetitive nature and suppressed recombination characteristic of pericentromeric regions in *D. melanogaster*.

#### Correlation tests

We calculated correlations between recombination rate and the estimated frequency of OreR at generation F25. Estimated F25 ancestry frequencies were averaged across all 15 replicates in non-overlapping 100 kb windows tiled across each chromosome arm using the Comeron et al. recombination map (FBrf0233439, [Comeron et al., 2012]; stdpopsim lift over to dm6 [Adrion et al., 2020]) as the reference coordinate system. Previous studies have also found a relationship between ancestry and the density of coding sites (CDS regions) [Schumer et al., 2018]; therefore, we also tested for these correlations. The number of CDS features overlapping each non-overlapping 100 kb window was obtained by interval overlap with *D. melanogaster* genome annotation (r6.57; Flybase). For the recombination rate [Comeron et al., 2012] and CDS features, we calculated the Spearman rank correlations and partial Spearman correlations (implemented in the ppcor R package) as recombination and CDS feature counts are positively correlated.

#### Identification of SNPs associated with selection

We tested every bi-allelic site on the autosomes and X chromosome for selection and/or epistatic selection associated with the mtDNA background by fitting the following quasibinomial GLM:

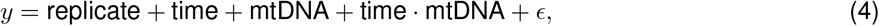

where *y* is the count of A and a alleles and *ϵ* is a binomially distributed error term. Then for each SNP, we calculated *p*-values for the effect of time from likelihood ratio tests with a reduced model:

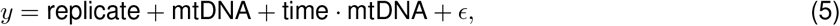

and repeated this for the mtDNA-by-time interaction effect:

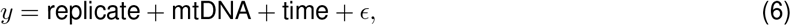

Model fitting and likelihood ratio tests were implemented using custom versions of the PoolFreqDiff scripts [Wiberg et al., 2017] using site coverage *>* 30 and *<* 200 filters.

## Acknowledgments

The work was supported by grants from NIGMS: MIRA R35GM139607 to D. M. R., L. J. D. was supported by NIH T32 GM128596 and an NSF Graduate Fellowship (2023-2025). D. M. R. acknowledges support of COBRE award P20GM109035.

## Author contributions

L.J.D. wrote the manuscript, conducted data analysis, and conceptualized the experiment. R.Z.B. maintained populations, conducted phenotype experiments, and contributed to the writing of the Methods section. F.A.L. conducted phenotype experiments and performed initial backcrosses. Y.R. conducted phenotype experiments, performed the round robin crosses, contributed data analysis pipelines, and conceptualized the experiment. D.M.R. edited the manuscript and conceptualized the experiment.

## Data Accessibility Statement

All DNA sequences have been deposited to NCBI (accession). All code and additional data needed to reproduce analysis and figures is available on GitHub (https://github.com/leahdarwin/RRpopCage).

## Conflict of Interest Statement

The authors declare no competing interests.

## Notes

### Competing Interest Statement

The authors have declared no competing interest.

## References

Adrion, J. R., Cole, C. B., Dukler, N., Galloway, J. G., Gladstein, A. L., Gower, G., Kyriazis, C. C., Ragsdale, A. P., Tsambos, G., Baumdicker, F., Carlson, J., Cartwright, R. A., Durvasula, A., Gronau, I., Kim, B. Y., McKenzie, P., Messer, P. W., Noskova, E., Ortega-Del Vecchyo, D., Racimo, F., Struck, T. J., Gravel, S., Gutenkunst, R. N., Lohmueller, K. E., Ralph, P. L., Schrider, D. R., Siepel, A., Kelleher, J., and Kern, A. D. (2020). A community-maintained standard library of population genetic models. eLife, 9:e54967.

Arguello, J. R., Laurent, S., and Clark, A. G. (2019). Demographic History of the Human Commensal Drosophila melanogaster. Genome Biology and Evolution, 11(3):844–854.

Bikard, D., Patel, D., Le Metté, C., Giorgi, V., Camilleri, C., Bennett, M. J., and Loudet, O. (2009). Divergent Evolution of Duplicate Genes Leads to Genetic Incompatibilities Within A. thaliana. Science, 323(5914):623–626.

Biot-Pelletier, D., Bettinazzi, S., Gagnon-Arsenault, I., Dubé, A. K., Bédard, C., Nguyen, T. H. M., Fiumera, H. L., Breton, S., and Landry, C. R. (2023). Evolutionary Trajectories are Contingent on Mitonuclear Interactions. Molecular Biology and Evolution, 40(4):msad061.

Burton, R. S. and Barreto, F. S. (2012). A disproportionate role for mtDNA in Dobzhansky–Muller incompatibilities? Molecular Ecology, 21(20):4942–4957.

Camus, M. F. and Inwongwan, S. (2023). Mitonuclear interactions modulate nutritional preference. Biology Letters, 19(12):20230375.

Chaturvedi, S., Lucas, L. K., Buerkle, C. A., Fordyce, J. A., Forister, M. L., Nice, C. C., and Gompert, Z. (2020). Recent hybrids recapitulate ancient hybrid outcomes. Nature Communications, 11(1):2179.

Chyb, S. and Gompel, N. (2013). Wild-type morphology. In Chyb, S. and Gompel, N., editors, Atlas of Drosophila Morphology, pages 1–23. Academic Press, San Diego.

Comeron, J. M., Ratnappan, R., and Bailin, S. (2012). The Many Landscapes of Recombination in Drosophila melanogaster. PLoS Genetics, 8(10):e1002905.

Darwin, L. J., Lemieux, F. A., Bachtel, R. Z., Blocker, J. H., Brown, C. P., Lerman, J. D., Maule, O. C., Raynes, Y., and Rand, D. M. (2025). Genetic and environmental interactions outweigh mitonu-clear coevolution for complex traits in Drosophila. ISSN: 2692-8205 Pages: 2025.11.24.689096 Section: New Results.

Duchen, P., Ž ivković, D., Hutter, S., Stephan, W., and Laurent, S. (2013). Demographic Inference Reveals African and European Admixture in the North American Drosophila melanogaster Population. Genetics, 193(1):291–301.

Farleigh, K., Highland, D. K., Alderman, M. G., Francioli, Y., Hirst, S. R., Faber, E. M., Perry, B. W., Holding, M. L., Castañeda-Gaytán, G., Borja, M., Franz-Chávez, H., Parkinson, C. L., Strickland, J. L., Margres, M. J., Mackessy, S. P., Meik, J. M., Castoe, T. A., and Schield, D. R. (2026). Evolution of genome-wide barriers to gene flow during complex speciation in rattlesnakes. Proceedings of the National Academy of Sciences, 123(21):e2609058123.

Gershoni, M., Templeton, A. R., and Mishmar, D. (2009). Mitochondrial bioenergetics as a major motive force of speciation. BioEssays, 31(6):642–650.

Grenier, J. K., Arguello, J. R., Moreira, M. C., Gottipati, S., Mohammed, J., Hackett, S. R., Boughton, R., Greenberg, A. J., and Clark, A. G. (2015). Global Diversity Lines–A Five-Continent Reference Panel of Sequenced Drosophila melanogaster Strains. G3 Genes|Genomes|Genetics, 5(4):593–603.

Hill, G. E. (2017). The mitonuclear compatibility species concept. The Auk, 134(2):393–409.

Huang, W., Massouras, A., Inoue, Y., Peiffer, J., Ràmia, M., Tarone, A. M., Turlapati, L., Zichner, T., Zhu, D., Lyman, R. F., Magwire, M. M., Blankenburg, K., Carbone, M. A., Chang, K., Ellis, L. L., Fernandez, S., Han, Y., Highnam, G., Hjelmen, C. E., Jack, J. R., Javaid, M., Jayaseelan, J., Kalra, D., Lee, S., Lewis, L., Munidasa, M., Ongeri, F., Patel, S., Perales, L., Perez, A., Pu, L., Rollmann, S. M., Ruth, R., Saada, N., Warner, C., Williams, A., Wu, Y.-Q., Yamamoto, A., Zhang, Y., Zhu, Y., Anholt, R. R. H., Korbel, J. O., Mittelman, D., Muzny, D. M., Gibbs, R. A., Barbadilla, A., Johnston, J. S., Stone, E. A., Richards, S., Deplancke, B., and Mackay, T. F. C. (2014). Natural variation in genome architecture among 205 Drosophila melanogaster Genetic Reference Panel lines. Genome Research, 24(7):1193–1208.

Immonen, E., Rö nn, J., Watson, C., Berger, D., and Arnqvist, G. (2016). Complex mitonuclear interactions and metabolic costs of mating in male seed beetles. Journal of Evolutionary Biology, 29(2):360–370.

Juric, I., Aeschbacher, S., and Coop, G. (2016). The Strength of Selection against Neanderthal Introgression. PLOS Genetics, 12(11):e1006340.

Kapun, M., Barrón, M. G., Staubach, F., Obbard, D. J., Wiberg, R. A. W., Vieira, J., Goubert, C., Rota-Stabelli, O., Kankare, M., Bogaerts-Márquez, M., Haudry, A., Waidele, L., Kozeretska, I., Pasyukova, E. G., Loeschcke, V., Pascual, M., Vieira, C. P., Serga, S., Montchamp-Moreau, C., Abbott, J., Gibert, P., Porcelli, D., Posnien, N., Sánchez-Gracia, A., Grath, S., Sucena, É., Bergland, A. O., Guerreiro, M. P. G., Onder, B. S., Argyridou, E., Guio, L., Schou, M. F., Deplancke, B., Vieira, C., Ritchie, M. G., Zwaan, B. J., Tauber, E., Orengo, D. J., Puerma, E., Aguadé, M., Schmidt, P., Parsch, J., Betancourt, A. J., Flatt, T., and González, J. (2020). Genomic Analysis of European Drosophila melanogaster Populations Reveals Longitudinal Structure, Continent-Wide Selection, and Previously Unknown DNA Viruses. Molecular Biology and Evolution, 37(9):2661–2678.

Kofler, R., Pandey, R. V., and Schlö \tterer, C. (2011). PoPoolation2: identifying differentiation between populations using sequencing of pooled DNA samples (Pool-Seq). Bioinformatics, 27(24):3435–3436.

Langdon, Q. K., Powell, D. L., Kim, B., Banerjee, S. M., Payne, C., Dodge, T. O., Moran, B., Fascinetto-Zago, P., and Schumer, M. (2022). Predictability and parallelism in the contemporary evolution of hybrid genomes. PLOS Genetics, 18(1):e1009914.

Li, H. and Durbin, R. (2009). Fast and accurate short read alignment with Burrows–Wheeler transform. Bioinformatics, 25(14):1754–1760.

Linder, R. A., Majumder, A., Chakraborty, M., and Long, A. (2020). Two Synthetic 18-Way Outcrossed Populations of Diploid Budding Yeast with Utility for Complex Trait Dissection. Genetics, 215(2):323–342.

Lopez, K. A., McDiarmid, C. S., Griffith, S. C., Lovette, I. J., and Hooper, D. M. (2021). Evaluating evidence of mitonuclear incompatibilities with the sex chromosomes in an avian hybrid zone. Evolution, 75(6):1395–1414.

Lynch, M. and Blanchard, J. L. (1998). Deleterious mutation accumulation in organelle genomes. Genetica, 102–103(1-6):29–39.

Lynch, M., Koskella, B., and Schaack, S. (2006). Mutation Pressure and the Evolution of Organelle Genomic Architecture. Science, 311(5768):1727–1730.

Ma, L., Clark, A. G., and Keinan, A. (2013). Gene-Based Testing of Interactions in Association Studies of Quantitative Traits. PLOS Genetics, 9(2):e1003321.

Maeda, G. P., Iannello, M., McConie, H. J., Ghiselli, F., and Havird, J. C. (2021). Relaxed selection on male mitochondrial genes in DUI bivalves eases the need for mitonuclear coevolution. Journal of Evolutionary Biology, 34(11):1722–1736.

Mallard, F., Nolte, V., Tobler, R., Kapun, M., and Schlötterer, C. (2018). A simple genetic basis of adaptation to a novel thermal environment results in complex metabolic rewiring in Drosophila. Genome Biology, 19(1):119.

Matute, D. R., Comeault, A. A., Earley, E., Serrato-Capuchina, A., Peede, D., Monroy-Eklund, A., Huang, W., Jones, C. D., Mackay, T. F. C., and Coyne, J. A. (2020). Rapid and Predictable Evolution of Admixed Populations Between Two Drosophila Species Pairs. Genetics, 214(1):211–230.

Meiklejohn, C. D., Holmbeck, M. A., Siddiq, M. A., Abt, D. N., Rand, D. M., and Montooth, K. L. (2013). An Incompatibility between a mitochondrial tRNA and its nuclear-encoded tRNA synthetase compromises development and fitness in Drosophila. PLoS genetics, 9(1):e1003238.

Moran, B. M., Payne, C., Langdon, Q., Powell, D. L., Brandvain, Y., and Schumer, M. (2021). The genomic consequences of hybridization. eLife, 10:e69016.

Moran, B. M., Payne, C. Y., Powell, D. L., Iverson, E. N. K., Donny, A. E., Banerjee, S. M., Langdon, Q. K., Gunn, T. R., Rodriguez-Soto, R. A., Madero, A., Baczenas, J. J., Kleczko, K. M., Liu, F., Matney, R., Singhal, K., Leib, R. D., Hernandez-Perez, O., Corbett-Detig, R., Frydman, J., Gifford, C., Schartl, M., Havird, J. C., and Schumer, M. (2024). A lethal mitonuclear incompatibility in complex I of natural hybrids. Nature, 626(7997):119–127.

Mossman, J. A., Biancani, L. M., Zhu, C.-T., and Rand, D. M. (2016). Mitonuclear Epistasis for Development Time and Its Modification by Diet in Drosophila. Genetics, 203(1):463–484.

Mossman, J. A., Ge, J. Y., Navarro, F., and Rand, D. M. (2019). Mitochondrial DNA Fitness Depends on Nuclear Genetic Background in Drosophila. G3 Genes |Genomes | Genetics, 9(4):1175–1188.

Nguyen, T. H. M., Klein, D. A., Weklar, O. S., Wengrow, E. R., and Rockman, M. V. (2025). Abundant Recurrent Mitochondrial Mutations and Widespread Mitonuclear Epistasis in Caenorhabditis elegans. Molecular Biology and Evolution, 42(12):msaf300.

Nguyen, T. H. M., Tinz-Burdick, A., Lenhardt, M., Geertz, M., Ramirez, F., Schwartz, M., Toledano, M., Bonney, B., Gaebler, B., Liu, W., Wolters, J. F., Chiu, K., Fiumera, A. C., and Fiumera, H. L. (2023). Mapping mitonuclear epistasis using a novel recombinant yeast population. PLOS Genetics, 19(3):e1010401.

Orr, H. A. and Turelli, M. (2001). THE EVOLUTION OF POSTZYGOTIC ISOLATION: ACCUMULATING DOBZHANSKY-MULLER INCOMPATIBILITIES. Evolution, 55(6):1085–1094.

Osada, N. and Akashi, H. (2012). Mitochondrial–Nuclear Interactions and Accelerated Compensatory Evolution: Evidence from the Primate Cytochrome c Oxidase Complex. Molecular Biology and Evolution, 29(1):337–346.

Pereira, R. J., Lima, T. G., Pierce-Ward, N. T., Chao, L., and Burton, R. S. (2021). Recovery from hybrid breakdown reveals a complex genetic architecture of mitonuclear incompatibilities. Molecular Ecology, 30(23):6403–6416. eprint: https://onlinelibrary.wiley.com/doi/pdf/10.1111/mec.15985.

Rand, D. M., Haney, R. A., and Fry, A. J. (2004). Cytonuclear coevolution: the genomics of cooperation. Trends in Ecology & Evolution, 19(12):645–653.

Robles, N. V., Moran, B. M., Rodríguez Barrera, M. J., Jofre, G. I., Gunn, T., Iverson, E. N. K., Beskid, S., Baczenas, J. J., Sedghifar, A., Andolfatto, P., Powell, D. L., Brandvain, Y., Havird, J. C., Rosenthal, G. G., and Schumer, M. (2025). Admixture Mapping Reveals Evidence for Multiple Mitonuclear Incompatibilities in Swordtail Fish Hybrids. Molecular Ecology, 34(22):e70106.

Sawyer, S. A. and Hartl, D. L. (1992). Population genetics of polymorphism and divergence. Genetics, 132(4):1161–1176.

Schlötterer, C. (2023). How predictable is adaptation from standing genetic variation? Experimental evolution in Drosophila highlights the central role of redundancy and linkage disequilibrium. Philosophical Transactions of the Royal Society B: Biological Sciences, 378(1877):20220046.

Schumer, M., Cui, R., Powell, D. L., Dresner, R., Rosenthal, G. G., and Andolfatto, P. (2014). High-resolution mapping reveals hundreds of genetic incompatibilities in hybridizing fish species. eLife, 3:e02535.

Schumer, M., Xu, C., Powell, D. L., Durvasula, A., Skov, L., Holland, C., Blazier, J. C., Sankararaman, S., Andolfatto, P., Rosenthal, G. G., and Przeworski, M. (2018). Natural selection interacts with recombination to shape the evolution of hybrid genomes. Science, 360(6389):656–660.

Sloan, D. B., Havird, J. C., and Sharbrough, J. (2017). The on-again, off-again relationship between mitochondrial genomes and species boundaries. Molecular Ecology, 26(8):2212–2236.

Spierer, A. N., Yoon, D., Zhu, C.-T., and Rand, D. M. (2021). FreeClimber: automated quantification of climbing performance in Drosophila. Journal of Experimental Biology, 224(2):jeb229377.

Tamura, K. (2003). Temporal Patterns of Fruit Fly (Drosophila) Evolution Revealed by Mutation Clocks. Molecular Biology and Evolution, 21(1):36–44.

Taylor, S. A. and Larson, E. L. (2019). Insights from genomes into the evolutionary importance and prevalence of hybridization in nature. Nature Ecology & Evolution, 3(2):170–177.

Van der Auwera, G. A., Carneiro, M. O., Hartl, C., Poplin, R., Del Angel, G., Levy-Moonshine, A., Jordan, T., Shakir, K., Roazen, D., Thibault, J., Banks, E., Garimella, K. V., Altshuler, D., Gabriel, S., and DePristo, M. A. (2013). From FastQ data to high confidence variant calls: the Genome Analysis Toolkit best practices pipeline. Current Protocols in Bioinformatics, 43(1110):11.10.1–11.10.33.

Weaver, R. J., Rabinowitz, S., Thueson, K., and Havird, J. C. (2022). Genomic Signatures of Mitonuclear Coevolution in Mammals. Molecular Biology and Evolution, 39(11):msac233.

White, M. A., Stubbings, M., Dumont, B. L., and Payseur, B. A. (2012). Genetics and Evolution of Hybrid Male Sterility in House Mice. Genetics, 191(3):917–934.

Wiberg, R. A. W., Gaggiotti, O. E., Morrissey, M. B., and Ritchie, M. G. (2017). Identifying consistent allele frequency differences in studies of stratified populations. Methods in Ecology and Evolution, 8(12):1899–1909.

Zuellig, M. P. and Sweigart, A. L. (2018). A two-locus hybrid incompatibility is widespread, polymorphic, and active in natural populations of Mimulus *. Evolution, 72(11):2394–2405.

