## Supplementary Figures and Tables for "Do divergent mitochondrial backgrounds alter the fate of nuclear alleles during hybridization and admixture?"

### Supplement

#### Figures

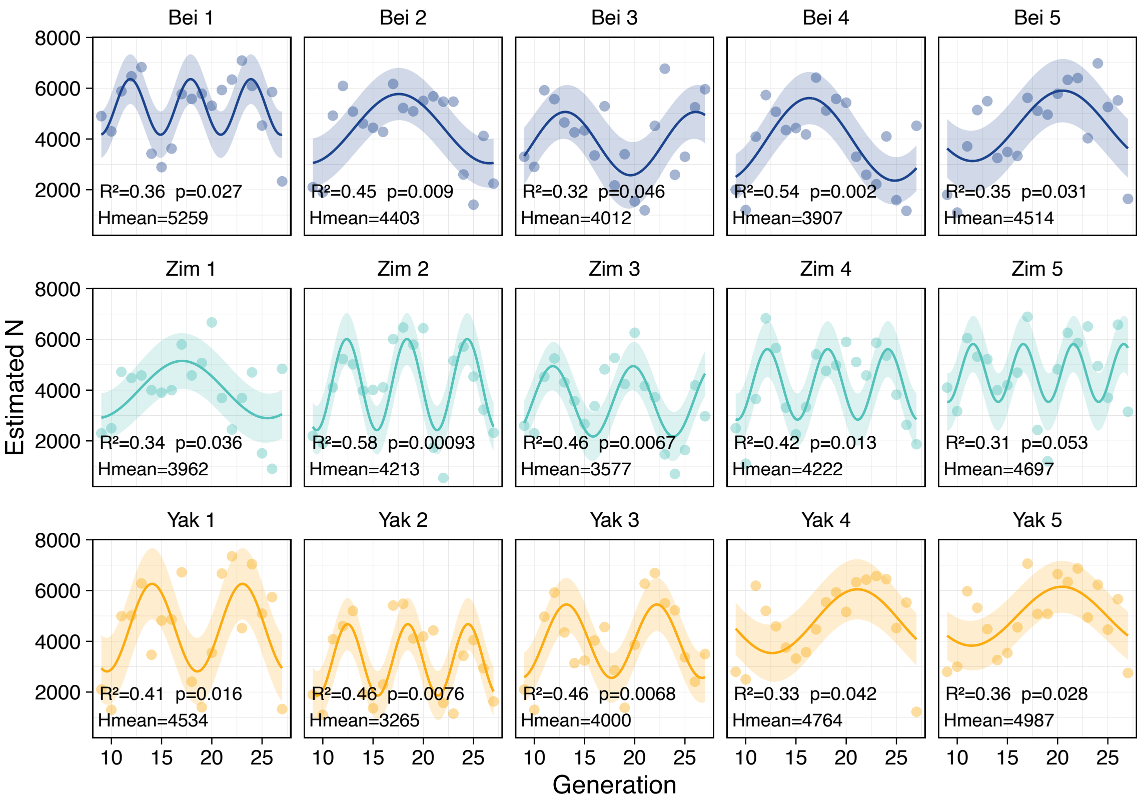


**Figure S1: Linear harmonic regressions of estimated population size for all populations.** For each population (mtDNA;replicate#), frozen adults were weighed between generations F7–F27 and used as a proxy for the census population size (points on plot). We observed fluctuations in population size likely related to larval density effects and fit linear harmonic regressions to the data (line on plot). The $R^2$ and $p$-value of the fit for each regression is given on the plot. Additionally, the census size over the total course of the experiment is estimated by the average of predicted $N$ values from the regression (Hmean).

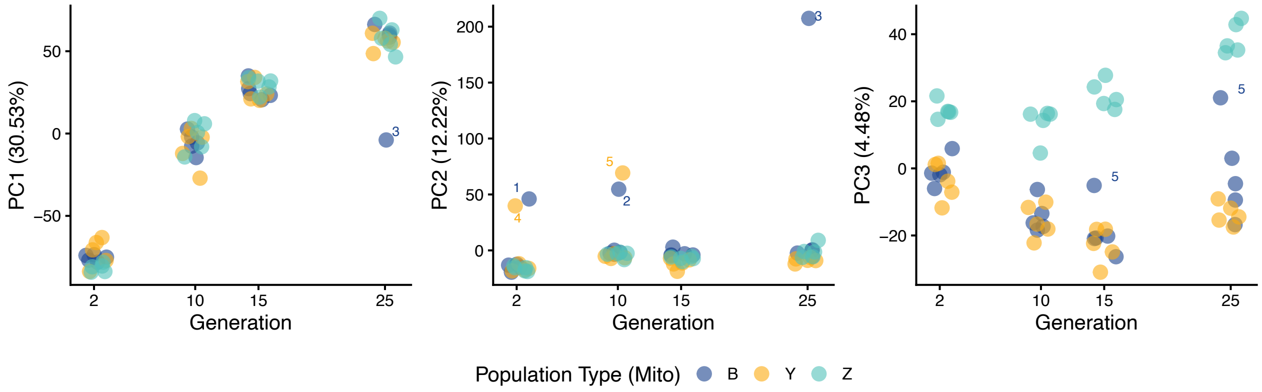


**Figure S2: Principal component (PC) decomposition of all allele frequencies over generation time.** Principal component 1 (PC1, left) over generations time explains 30.53% in genetic variation and increases with generation time across all mtDNA (Mito) population types. Principal component 2 (PC2, middle) explains 12.22% of genetic variation and is attributable to specific population outliers (replicate number labeled near points). Principal component 3 (PC3, right) explains 4.48% of genetic variation and shows a divergence in the populations with the Zimbabwe (Z) mtDNA relative to Beijing (B) and *D. yakuba* mtDNA populations over time with the exception of Beijing replicate 5.

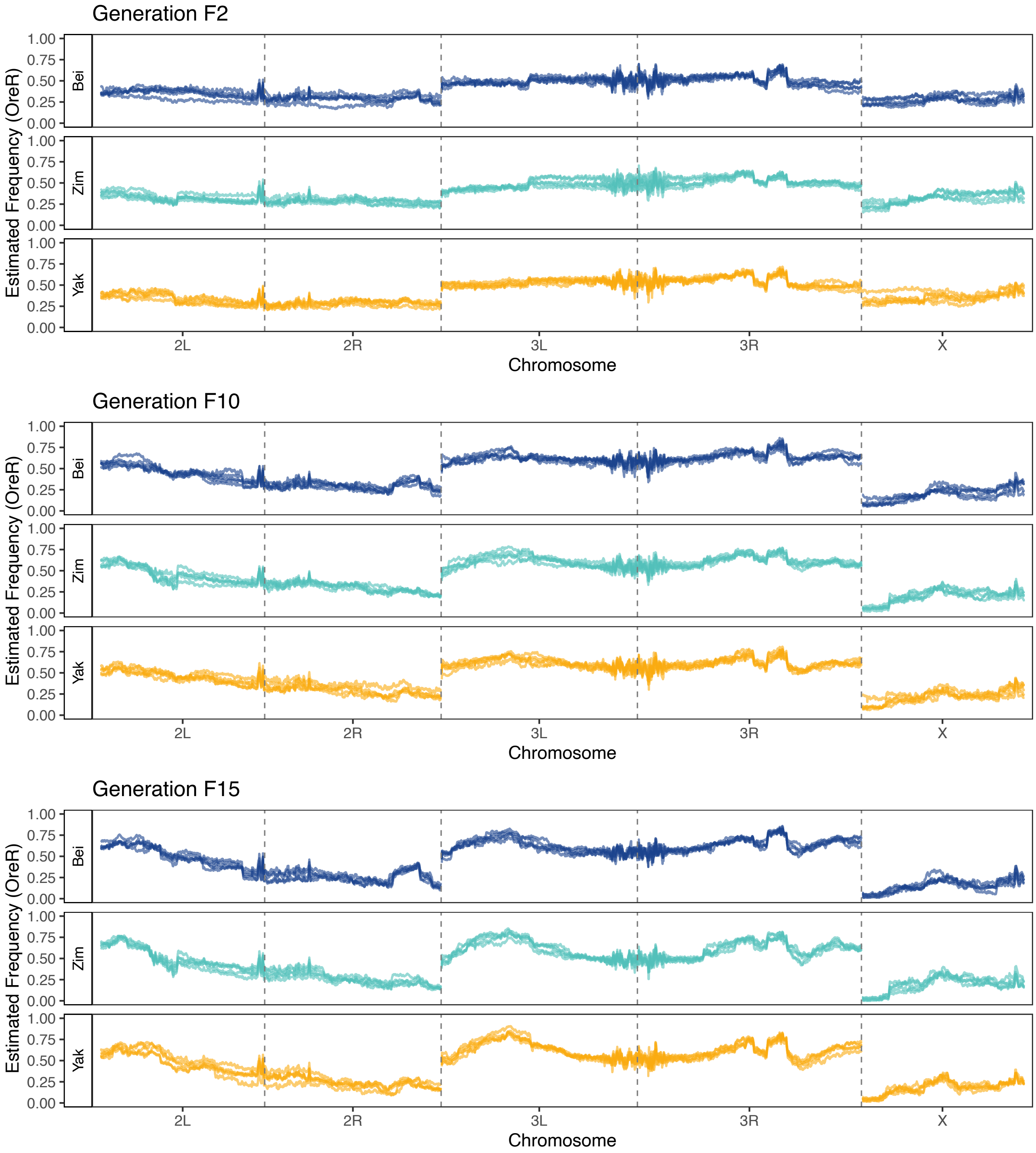


**Figure S3: Estimated ancestry frequencies for generations F2, F10, F15.** Plots show estimated frequencies of the OreR ancestry for all replicate populations. Populations are colored by myDNA type Beijing (Bei), Zimbabwe (Zim), and *D. yakuba* (Yak). The Ral375 ancestry is not plotted but note that it is at frequency 1-OreR.

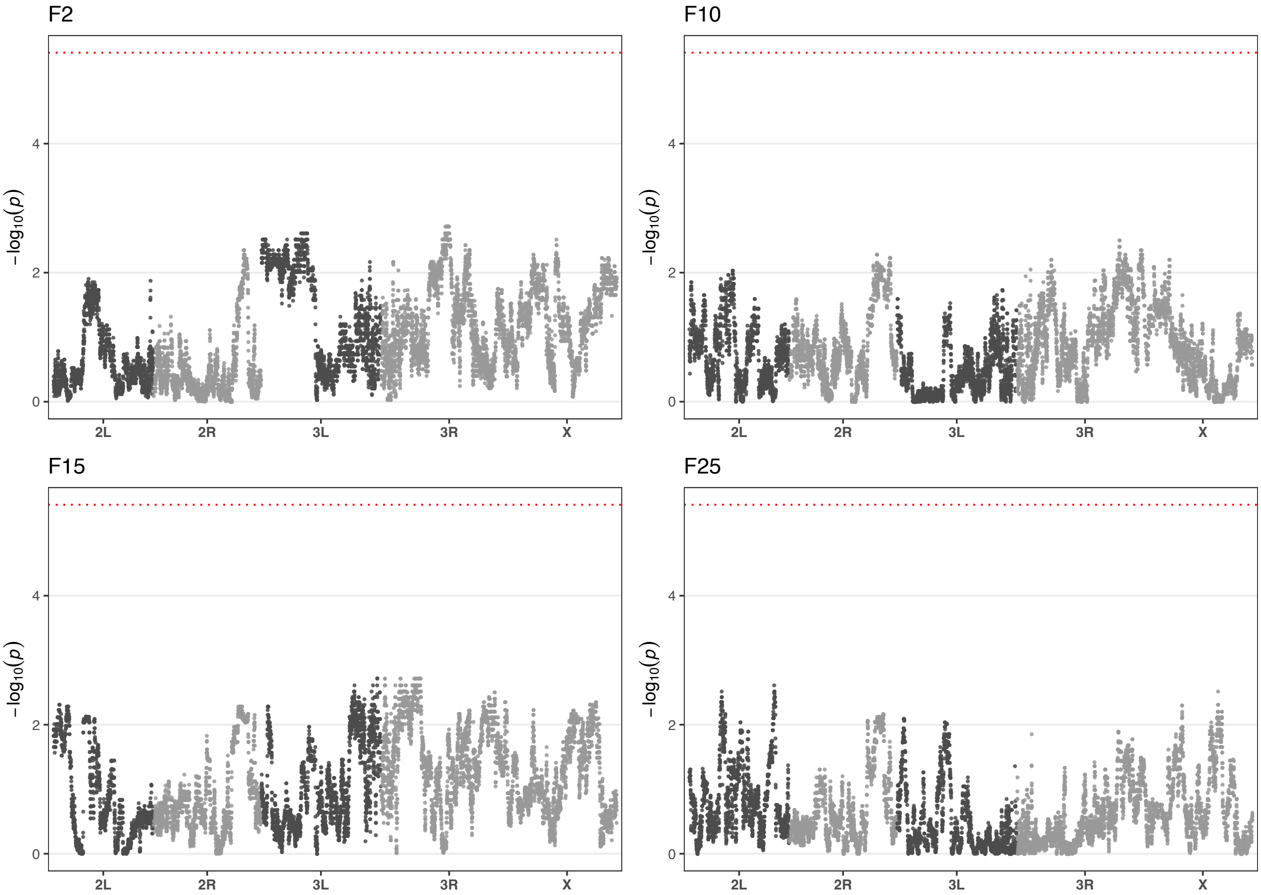


**Figure S4: Windowed Kruskal-Wallis test for mtDNA effects on ancestry proportion for all sampled generations.** A manhattan plot of $p$-values windowed Kruskal-Wallis (KW) test at each generation (F2, F10, F15, F25) is shown. A significant KW test (above the Bonferroni corrected dotted red line) indicates an mtDNA effect on the ancestry proportion in the given window.

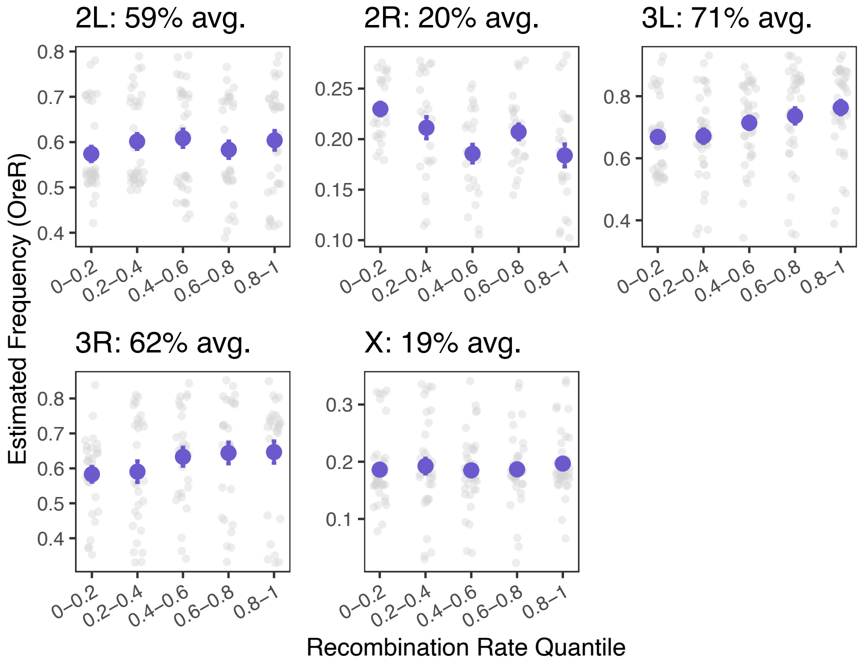


**Figure S5: Correlations between OreR ancestry and recombination rate.** For each chromosome arm, the relationship between the estimated OreR ancestry and recombination rate ( [Comeron et al., 2012]) is shown. In each title, the total proportion of OreR for the chromosome arm is given. Recombination rates are grouped into five quantiles on the x-axis for visualization and the mean ancestry for each quantile is plotted in purple with standard error bars. Correlation coefficients and $p$-values are given in Table S3.

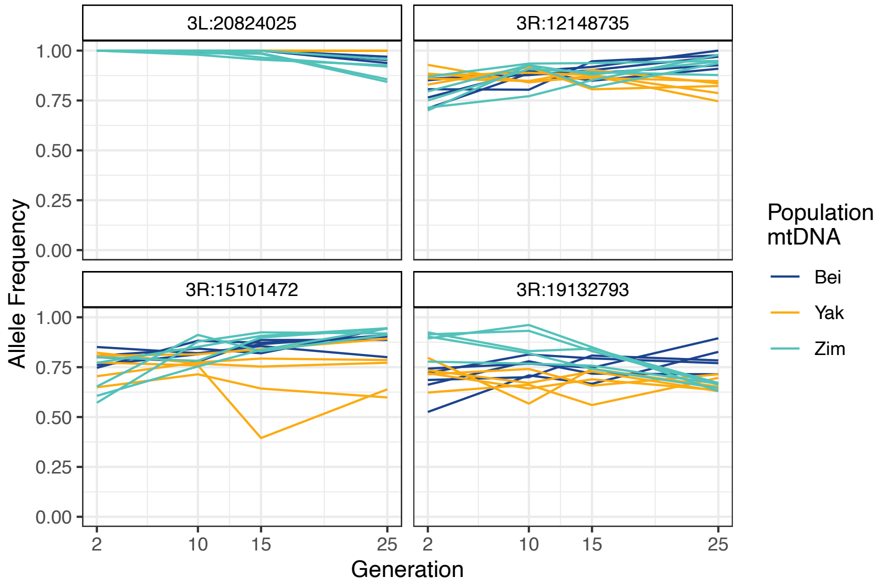


**Figure S6: Allele frequency trajectories for significant SNPs.** For SNPs (MAF>0.01) not associated with the $w^{1118}$ background but with a significant mtDNA-by-time interaction effect per likelihood ratio test, the allele frequency trajectories are plotted. Trajectories are colored by mtDNA background: Beijing (Bei), Zimbabwe (Zim), and *D. yakuba* (Yak). The panel label gives the chromosome arm and position of the allele.

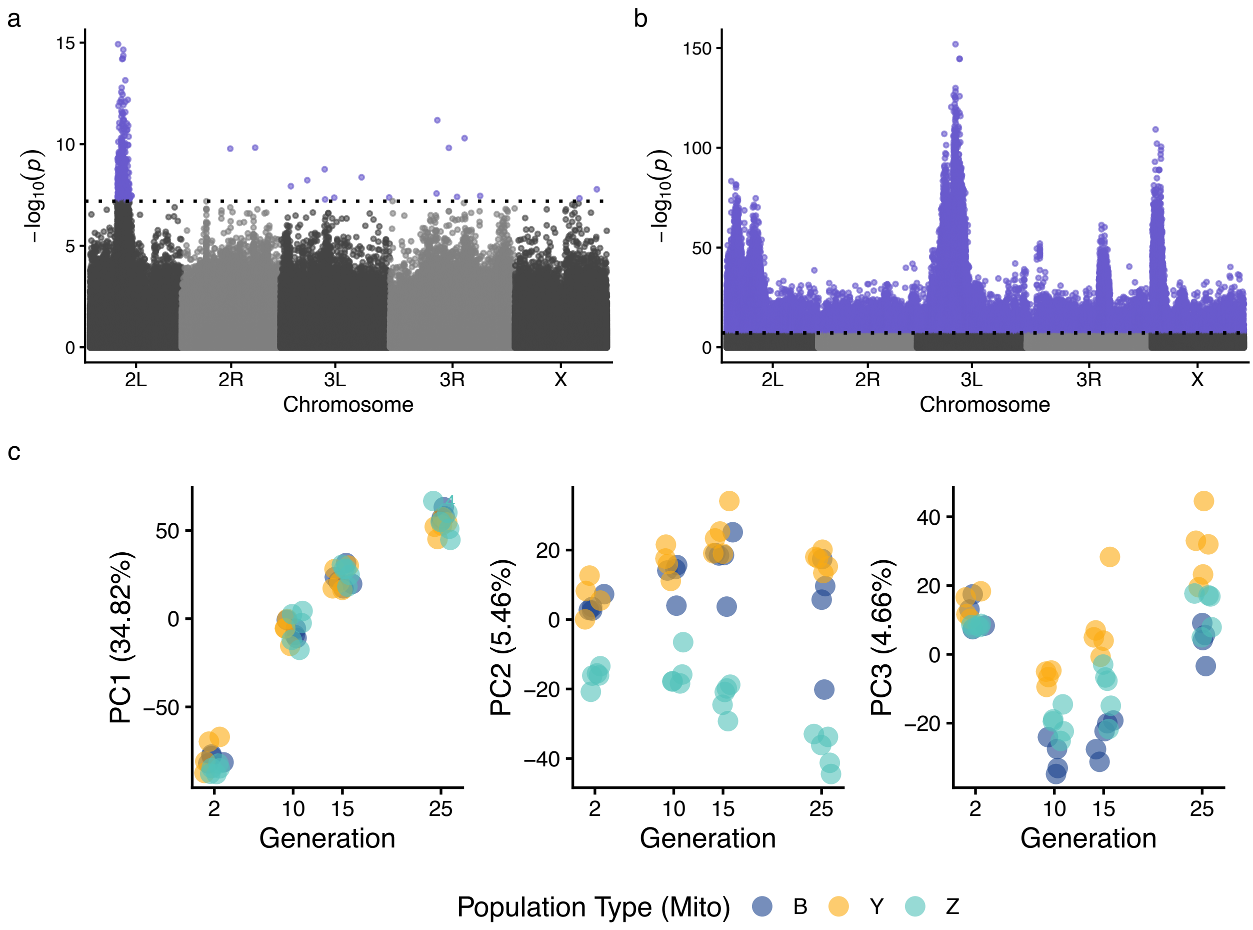


**Figure S7: Data reanalysis excluding PCA outliers.** The following outliers were excluded prior to reanalysis and are shown in Figure S2 (Mito;Replicate#;Generation): Bei;3;F25, Bei;1;F2, Bei;2;F10, Yak;4;F2, Yak;5;F10. **(a)** The manhattan plot shows the reanalysis of Figure 4a of the $p$-values for likelihood ratio test on time×mtDNA interaction effect. **(b)** The manhattan plot shows the reanalysis of Figure 3a of the $p$-values for likelihood ratio test on time effect. The dotted lines in panels (a) and (b) show the Bonferroni-adjusted significance threshold and highlighted points are significant. **(c)** The three plots show a reanalysis of PC1–3 over generation time from Figure S2. Points are colored by population type for each mtDNA background Beijing (B), *D. yakuba* (Y), and Zimbabwe (Z).

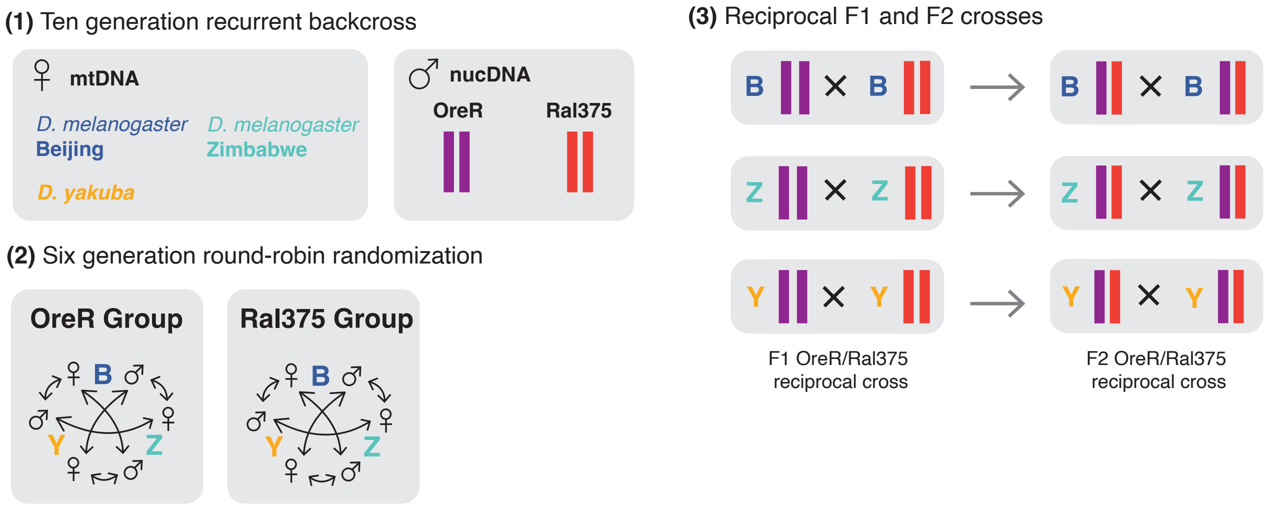


**Figure S8: Extended methods for experimental evolution of hybrids.** The experimental hybrid populations were initiated in three distinct steps with the goal of creating similar nuclear hybrid backgrounds for each of the mtDNA haplotypes. First, the three mtDNA haplotypes were recurrently backcrossed onto two nuclear backgrounds (nucDNA) over ten generations. Second, within each backcross group (OreR/Ral375), the nuclear backgrounds were randomized across mtDNA types using a six generation round-robin crossing scheme. Third, following the round-robin, reciprocal F1 and F2 crosses were using to create hybrid nuclear genomes within each mtDNA background.

### Tables

**Table S1:** Mixed model ANOVA results for development time and fecundity.

| Trait | Term | Sum Sq | Mean Sq | Num DF | Den DF | $F$ | $p$ |
|---|---|---|---|---|---|---|---|
| **Development Time** | Mito | 308.072 | 154.036 | 2 | 6.043 | 1.874 | 0.233 |
| **Fecundity** | Mito | 215.800 | 107.900 | 2 | 12.000 | 0.283 | 0.759 |

**Table S2:** Estimated marginal means by trait and mitotype.

| Trait | Mitotype | Estimate | SE | Lower 95% CI | Upper 95% CI |
|---|---|---|---|---|---|
| Development Time | Bei | 108.034 | 0.960 | 105.670 | 110.398 |
| | Zim | 107.861 | 0.965 | 105.497 | 110.226 |
| | Yak | 110.243 | 0.974 | 107.877 | 112.609 |
| Fecundity | Bei | 46.591 | 3.726 | 38.474 | 54.709 |
| | Zim | 45.360 | 3.726 | 37.243 | 53.477 |
| | Yak | 49.236 | 3.726 | 41.118 | 57.353 |

**Table S3:** Spearman rank correlations between OreR frequency and recombination rate or CDS density. Partial correlations control for the other covariate.

| Chr | $\rho$ [95% CI] | $p$ | $\rho_{\text{partial}}$ [95% CI] | $p_{\text{partial}}$ |
|---|---|---|---|---|
| *Recombination rate* | | | | |
| All | -0.038 [-0.116, 0.041] | 0.350 | -0.031 [-0.110, 0.047] | 0.435 |
| 2L | 0.026 [-0.125, 0.177] | 0.734 | 0.017 [-0.135, 0.168] | 0.831 |
| 2R | -0.272 [-0.431, -0.097] | 0.003 | -0.255 [-0.416, -0.077] | 0.005 |
| 3L | 0.333 [0.200, 0.453] | 2.43e-06 | 0.332 [0.199, 0.452] | 2.80e-06 |
| 3R | 0.209 [0.046, 0.361] | 0.013 | 0.219 [0.055, 0.370] | 0.009 |
| X | 0.063 [-0.082, 0.205] | 0.397 | 0.056 [-0.089, 0.199] | 0.445 |
| *CDS density* | | | | |
| All | -0.063 [-0.141, 0.016] | 0.118 | -0.059 [-0.137, 0.019] | 0.140 |
| 2L | 0.081 [-0.071, 0.229] | 0.298 | 0.078 [-0.074, 0.227] | 0.315 |
| 2R | -0.148 [-0.320, 0.033] | 0.108 | -0.111 [-0.286, 0.071] | 0.232 |
| 3L | 0.055 [-0.087, 0.195] | 0.450 | 0.047 [-0.096, 0.188] | 0.518 |
| 3R | -0.059 [-0.222, 0.107] | 0.485 | -0.088 [-0.249, 0.079] | 0.301 |
| X | 0.104 [-0.040, 0.244] | 0.157 | 0.101 [-0.044, 0.241] | 0.172 |

**Table S4: First three generations of round robin crosses.** Listed are the directions of the crosses for each generation of the round robin crosses to homogenize the nuclear backgrounds. Line names are given as *mtDNA;nucDNA*. Subscripts on the nuclear genotype indicate the mtDNA backcross associated with the line and indicates that there was potentially private variation associated with that line confounding the mtDNA type. In the table, nuclear backgrounds become a mixture of the two parents and for simplicity are replaced with letter labels each generation. Only the Ral375 (Ral) group is shown but the process was identical for the OreR group. This process was repeated another 3 times, but generations F4–F6 are not given in the table for brevity.

| Virgin Females | Males | Offspring |
|---|---|---|
| **Generation F1** | | |
| ZW142;$Ral_{ZW142}$ | Bei11;$Ral_{Bei11}$ | ZW142;A |
| ZW142;$Ral_{ZW142}$ | Yak;$Ral_{Yak}$ | ZW142;B |
| Bei11;$Ral_{Bei11}$ | ZW142;$Ral_{ZW142}$ | Bei11; C |
| Bei11;$Ral_{Bei11}$ | Yak;$Ral_{Yak}$ | Bei11;D |
| Yak;$Ral_{Yak}$ | Bei11;$Ral_{Beii}$ | Yak;E |
| Yak;$Ral_{Yak}$ | ZW142;$Ral_{ZW142}$ | Yak;F |
| **Generation F2** | | |
| ZW142;A | Yak;F | ZW142;G |
| ZW142;B | Bei11;D | ZW142;H |
| Bei11; C | Yak;E | Bei11;I |
| Bei11;D | ZW142;B | Bei11;J |
| Yak;E | Bei11; C | Yak;K |
| Yak;F | ZW142;A | Yak;L |
| **Generation F3** | | |
| ZW142;G | Bei11;I | ... |
| ZW142;H | Yak;K | ... |
| Bei11;I | ZW142;G | ... |
| Bei11;J | Yak;L | ... |
| Yak;K | ZW142;H | ... |
| Yak;L | Bei11;J | ... |
